# Effects of theta transcranial alternating current stimulation (tACS) on exploration and exploitation during uncertain decision-making

**DOI:** 10.1101/2021.07.26.453851

**Authors:** Miles Wischnewski, Boukje Compen

## Abstract

Exploring ones surroundings may yield unexpected rewards, but is associated with uncertainty and risk. Alternatively, exploitation of certain outcomes is related to low risk, yet potentially better outcomes remain unexamined. As such, risk-taking behavior depends on perceived uncertainty and a trade-off between exploration-exploitation. Previously, it has been suggested that risk-taking may relate to theta activity in the prefrontal cortex. Furthermore, previous studies hinted at a relationship between a right-hemispheric bias in frontal theta asymmetry and risky behavior. In the present double-blind sham-controlled within-subject study, we applied bifrontal transcranial alternating current stimulation (tACS) at the theta frequency (5 Hz) on eighteen healthy volunteers during a gambling task. Two tACS montages with either left-right or posterior-anterior current flow were employed at an intensity of 1 mA. Results showed that, compared to sham, theta tACS increased perceived uncertainty irrespective of current flow direction. Despite this observation, no direct effect of tACS on exploration behavior and general risk-taking was observed. Furthermore, frontal theta asymmetry was more right-hemispherically biased after posterior-anterior tACS, compared to sham. Finally, we used electric field simulation to identify which regions were targeted by the tACS montages as an overlap in regions may explain why the two montages resulted in comparable outcomes. Our findings provide a first step towards understanding the relationship between frontal theta oscillations and different features of risk-taking.

**Highlights:**

– Risk taking is related to uncertainty, exploration and exploitation
– Frontal theta tACS was applied to modulate aspects of risk taking
– tACS did increase perceived uncertainty, but not exploration behavior
– tACS induced a right hemispheric shift in frontal theta asymmetry

## Introduction

In survival situations rewards are often associated with risk. For example, during foraging an agent is exposed to potential predators meaning that prolonged reward seeking behavior is coupled to increased risk. In humans, foraging-like behavior is observed in sequential decision making during uncertain situations, for example in gambling or stock trading. In these situations one has to constantly evaluate and update beliefs on a balance between risk and reward. This balancing act has been described as the exploration-exploitation trade-off, which poses that an agent seeks a compromise between exploiting rewards based on existing knowledge and exploring other options to gain knowledge of unknown rewards and losses [1-3]. As such, exploitation and exploration are akin to risk aversive and risk seeking behavior [1, 2].

The exploration-exploitation trade-off hypothesis suggests that decision-making during risky situations can be described in three non-mutually exclusive dimensions [3]. First, risk-taking depends on behavioral patterns, where more diverse actions relate to exploration and stable behavior relates to exploitation [4, 5]. Second, risk-taking is subject to the amount of uncertainty in a given situation [6]. While uncertainty often leads to a decrease in exploratory behavior due to general risk-aversive tendencies, it may promote exploration in predictable situations or when the current reward is unsatisfactory [6]. Third, risk-taking depends on the utility of the obtained outcome. Although exploration potentially increases the probability of a loss in one trial, the agent may obtain information on optimizing future behavior [1].

Besides different elemental features of risk-taking, various studies have investigated the neural correlates of risk-related decision-making [7-9]. Next to sub-cortical reward structures, such as the sub-thalamic nucleus and striatum, various areas in the prefrontal cortex (PFC) are involved in decision making under uncertainty [8-10]. Meder et al. [8] used a sequential decision making task in which accumulation of foraged reward was related to increased risk of losing all gains. It was found that hemodynamic activity in the dorsolateral PFC (DLPFC) and inferior frontal gyrus (IFG), among other areas, are linearly scaled with increased reward and risk probability. Furthermore, the DLPFC has been associated with top-down processes guiding exploration and risk seeking [7, 9-12]. Additionally, it has been suggested that reward-related areas such as the anterior cingulate cortex, orbitofrontal cortex and IFG drive exploitation of safe rewards [10, 13-15]. Indeed, IFG has been related to inhibitory control, which may, in turn, promote risk aversion [14, 16].

Electrophysiological recordings have shown that frontal cortex activity related to risk-taking is associated with oscillatory theta rhythms [17-20]. More specifically, frontal theta activation has been associated with reinforcement feedback processing, which is crucial for updating prediction models on future reward and punishment [17, 20-26]. For instance, Cavanagh et al. [17] showed that frontal theta oscillations are positively related to reward uncertainty, primarily in individuals in which uncertainty drives exploratory behavior. This may suggest that frontal theta-related prediction error processes are involved in updating reward/punishment contingencies during exploration [17].

The association between frontal theta oscillations and risk-taking behavior has been further explored by attempting to modulate decision-making under risk using transcranial alternating current stimulation (tACS) [19, 27-30]. TACS is thought to modulate endogenous oscillatory activity by applying exogenous sinusoidal currents to the scalp [31, 32]. On the one hand, Sela et al. [19] found that right frontal theta tACS may increase risk-related behavior. On the other hand, Dantas et al. [27] found decreased risk-taking when theta tACS was applied over the left frontal cortex. Together, these results may hint at frontal theta asymmetry as an underlying functional correlate of risk-related behavior, where increased risk-taking is associated with right frontally biased theta activity. Indeed, a number of imaging studies pointed to a relationship between risk-taking and an imbalance in frontal cortex activity [7, 16, 33], particularly for theta oscillations [18, 20]. However, results have been mixed and other studies have found no evidence for frontal asymmetry as a correlate for risk-taking [8, 34, 35], nor any effects of tACS [30]. For instance, Yaple et al. [30] found that neither left nor right lateralized frontal theta tACS affected risky behavior. Furthermore, Dantas et al. [27] reported that tACS-induced changes in risk-taking were not related to frontal theta asymmetry.

Since risk-taking behavior underlies complex multi-faceted cognitive processes, it is unsurprising that evidence on the efficacy of theta tACS has been mixed. In the present study, we therefore investigated the effect of theta-tACS on the exploration-exploitation dimensions ‘behavioral pattern’, ‘uncertainty’, and ‘value of obtained outcome’, related risk-related decision-making. Given the tentative evidence that frontal theta asymmetry potentially relates to risk-taking, two tACS montages were employed to target the PFC. In one montage, inter-hemispheric left-right current flow was applied. In this condition, tACS-induced currents in the left and right PFC are in opposing phase, which may promote hemispheric asynchrony, and consequently, theta asymmetry [36, 37]. In the other montage, an intra-hemispheric posterior-anterior current flow was applied. Since in this montage induced current is similar in both left and right PFC, we expected that this setup would promote synchronization, and consequently, hemispheric balance [36, 37]. Based on the established link between theta and uncertainty [17], we hypothesized that frontal theta tACS would alter subjective uncertainty, which could potentially yield adjustments in exploratory and general risk-taking behavior. Furthermore, we expected that these effects would be predominantly observed when tACS induces frontal theta asymmetry. Finally, to explore similarities between the two tACS montages, induced electric field distributions were compared and overlaid.

## Methods

### Participants

A total of 18 participants (mean ± SEM, 21.9 ± 2.3, 15 females) were included in the present study. Approval for the study was received from the medical ethical committee of Radboud University Medical Centre in Nijmegen, and participants were recruited using SONA, Radboud University’s online recruitment system. Participants were required to be between 18-35 years of age and right-handed. Exclusion criteria were use of psychotropic medication, head trauma or previous brain surgery, neurological or psychiatric disorders, large or ferromagnetic metal parts in the head, an implanted cardiac pacemaker or neurostimulator, pregnancy and skin disease. Furthermore, participants were excluded if they had participated in a non-invasive brain stimulation (NIBS) study less than 24 hours before.

### Risk taking task

A modified version of the sequential gambling task by Meder et al., (2016) was used, which induces a trade-off between accumulating reward and increasing risk. The task was programmed using Presentation software (version 17.1, www.neurobs.com). Participants were seated at a distance of approximately 80 cm from the screen on which the task was presented (50 × 30 cm, 1680 × 1050 pixels). A six-sided, white dice with red pips (380 × 380 pixels) was presented on a green background. Each trial started with a rolling phase, jittered between 1.5 and 3.0 s, during which one of the six random chosen sides of the dice was presented for 150 ms, and then replaced by another side. After the rolling phase, the randomly chosen outcome of the dice was presented for 2 s, together with the amount of points collected in the trial so far.

If the side of the dice with 1 pip was presented, no points were received and the trial ended. If one of the sides with 2, 3, 4, 5 or 6 pips was presented, participants received an amount of points corresponding to the number of pips, multiplied by ten. Participants were then provided with a choice between two actions. First, they could decide to continue and roll the dice again. Second, they could stop and bank the amount of points collected in the trial. These points were then added to the total score, after which the trial ended. The action of choice was indicated by pressing the left or right arrow key with the index fingers, and the association between actions and keys was counterbalanced across the participants. When the participant chose to continue rolling the dice, a new gambling round started after 2 seconds. Any outcomes higher than 1 pip resulted in points being added to the current balance, and the choice between continuing and stopping was then again presented. However, when presented with the side of the dice with 1 pip, the balance was set to 0, such that all points of that trial were lost, and the trial ended. When participants chose to stop, or when the outcome of a dice roll was 1 pip, they were presented with the amount of points earned in that trial for 2.5 seconds. At the end of each trial, the total score was shown for 3.5 seconds. A graphical representation of the experimental task is displayed in Figure 1.

**Figure 1.**
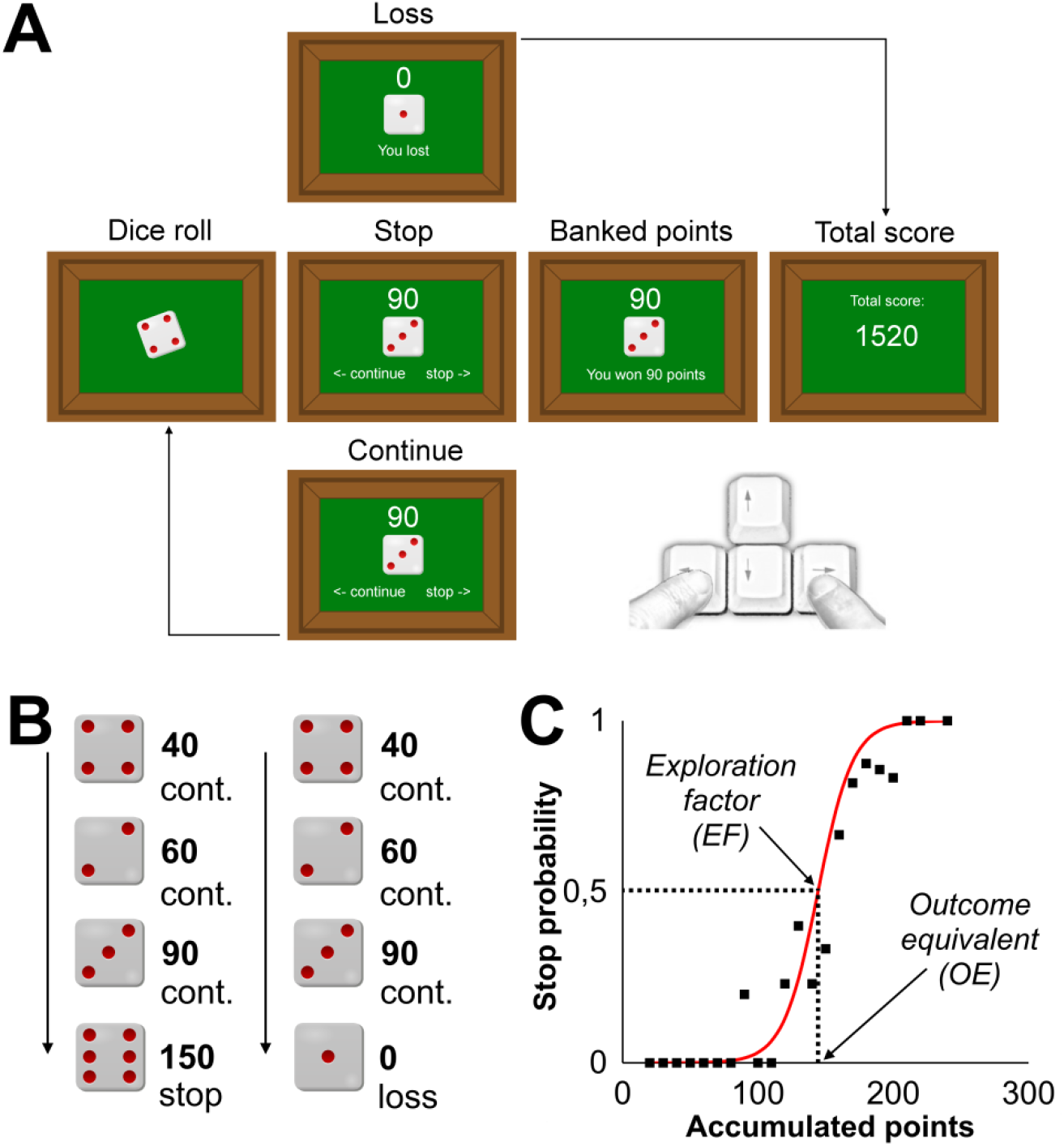
Sequential gambling task. A) Each trial started with a rolling phase (1.5 – 3.0 s), after which the dice stopped randomly at pips 1-6. The dice was displayed together with the number of summed points within the trials so far. Three possible scenarios follow: ‘Loss’ when the dice resulted in pip 1, which meant that all summed points were lost. ‘Continue’ when the dice resulted in pips 2-6 and participant chooses to continue for another round. ‘Stop’ when the dice resulted in pips 2-6 and participant decides to stop and banks the points collected in the trial, which are added to the total score. After stopping or losing the trial ends displaying the total score (3.5 seconds) and the next trial starts. B) An example trial containing four rounds where the participant continues (cont.) and then either stops at the fourth round (left), banking 150 points, or alternatively, the participant loses at the fourth round (right), gaining no points in this trial. C) The stop probability, which refers to the likelihood that a participant stops at a certain amount of summed points, is fitted by a 2-parameters logistic curve. The half-max and growth rate parameters of the function represent the outcome equivalent (OE) and exploration factor (EF), respectively. Together with the uncertainty value (UV), which is gathered from reaction times, these parameters reflect exploration-exploitation dimensions of decision-making in a risk environment.

Each session consisted of six blocks of 20 trials. Participants were asked to gain as many points as possible. Furthermore, they were required to make their choices within 2 seconds, otherwise the trial ended with 0 points.

### Behavioral outcomes

Choice (continue, stop, loss) and reaction time in each round within a trial (continue, stop, loss) were recorded. For each potential outcome we calculated the stop probability, which corresponds to the ratio between continue and stop choices. The distribution of stop probability (stop|x_n_) can be reliably modelled using a logistic function:

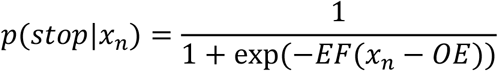

X_n_ reflects the accumulated points in a given trial n. The exploration factor (EF) corresponds to the growth rate of the logistic function. This free parameter is an estimation of the ‘behavioral pattern’ dimension with the exploration-exploitation framework and reflects the variability/stability of choices [3]. Larger values indicate a steeper curve, which reflect more stable exploitation behavior, whereas smaller values relate to exploratory behavior. The outcome equivalent (OE) reflects the inflection point of the logistic curve, where the probability of continuing and stopping is equal. The OE is a composite measure of the ‘behavioral patterns’ and ‘obtained outcome’ dimensions within the exploration-exploitation framework [3]. Larger OE is indicative of higher general risk taking and willingness to explore a high risk environment, which can provide information for future trials, at the cost of potential losses.

In addition to the chosen action, participants’ reaction times in each round of a trial were obtained. Average reaction times were calculated for the first round and last round where participants chose to continue. With an increasing number of rounds within a trial, and thus more accumulated points, participants’ uncertainty regarding whether or not to continue escalates. Therefore, last round reaction times, compared to first round reaction times, provide a measure for subjective uncertainty. As such, the uncertainty value (UV) was calculated, which reflects the difference between last and first round reaction times. The UV corresponds to the ‘uncertainty’ dimension of the exploration-exploitation framework, where higher values reflect more perceived uncertainty.

### EEG pre-processing and data reduction

EEG was recorded with the ActiveTwo system (BioSemi, Amsterdam, The Netherlands), using 32 active electrodes that were positioned according to the 10-20 system. An online 0.16 -100 Hz band-pass filter with a sampling rate of 2048 Hz was used. The common mode sense (CMS) and driven right leg (DRL) electrodes were placed over C1 and C2 and served as the online reference and ground electrodes. Resting-state EEG was recorded for four minutes, alternating every 60 seconds between eyes open and closed, and measured before and after stimulation.

The pre-and post-stimulation resting-state EEG was analysed using BrainVision Analyzer 2.1 (Brain Products GmbH, München, Germany). Raw EEG signal was re-referenced to the average signal of the 32 electrodes. EEG was segmented in epochs of two seconds, after which the Gratton and Coles method was employed to correct for vertical eye movements, using electrode Fp1 [38]. Additional artifacts were removed semi-automatically: signals that led to amplitudes -/+100 µV were marked, after which the data was visually inspected. An offline band-pass of 1 – 30 Hz (48 dB/octave) was applied. Spectral data was acquired by using a fast Fourier transformation (Hanning window length: 10%), which was averaged over all epochs in the four-minute resting-state EEG period. Spectral activity was calculated for the delta (1-4 Hz), theta (4 – 7 Hz), alpha (7-13 Hz), and beta (13-30 Hz) bands. Furthermore, frontal theta asymmetry was calculated by dividing left and right spectral activity divided by the total: F4 – F3 / F4 + F3 [39].

### Transcranial alternating current stimulation

Stimulation was delivered by a NeuroConn DC current stimulator (NeuroConn GmbH, Ilmenau, Germany). Four stimulation electrodes (5 × 3 cm) were placed to target the left and right prefrontal cortex, 2 cm lateral of AF3 and AF4 and 1 cm lateral of Fc1 and Fc2 [29]. This montage allowed for either intra-hemispheric (tACS_INTRA_) or inter-hemispheric (tACS_TRANS_) current flow (Figure 2) [36, 37]. Specifically, for tACS_TRANS_, polarity of AF3 and Fc1 electrodes was the same, as opposed to AF4 and Fc2 electrodes, yielding alternating left-right current flow (Figure 2A,B). For the tACS_INTRA_ condition, polarity of AF3 and AF4 electrodes was the same, as opposed to Fc1 and Fc2 electrodes, yielding alternating anterior-posterior current flow (Figure 2D,E). Although both montages target the PFC, the specific distribution of induced currents differs (Figure 2C,F). Conductive rubber electrodes were placed on the scalp under an EEG cap using adhesive Ten20 paste (Weaver and Company).

**Figure 2.**
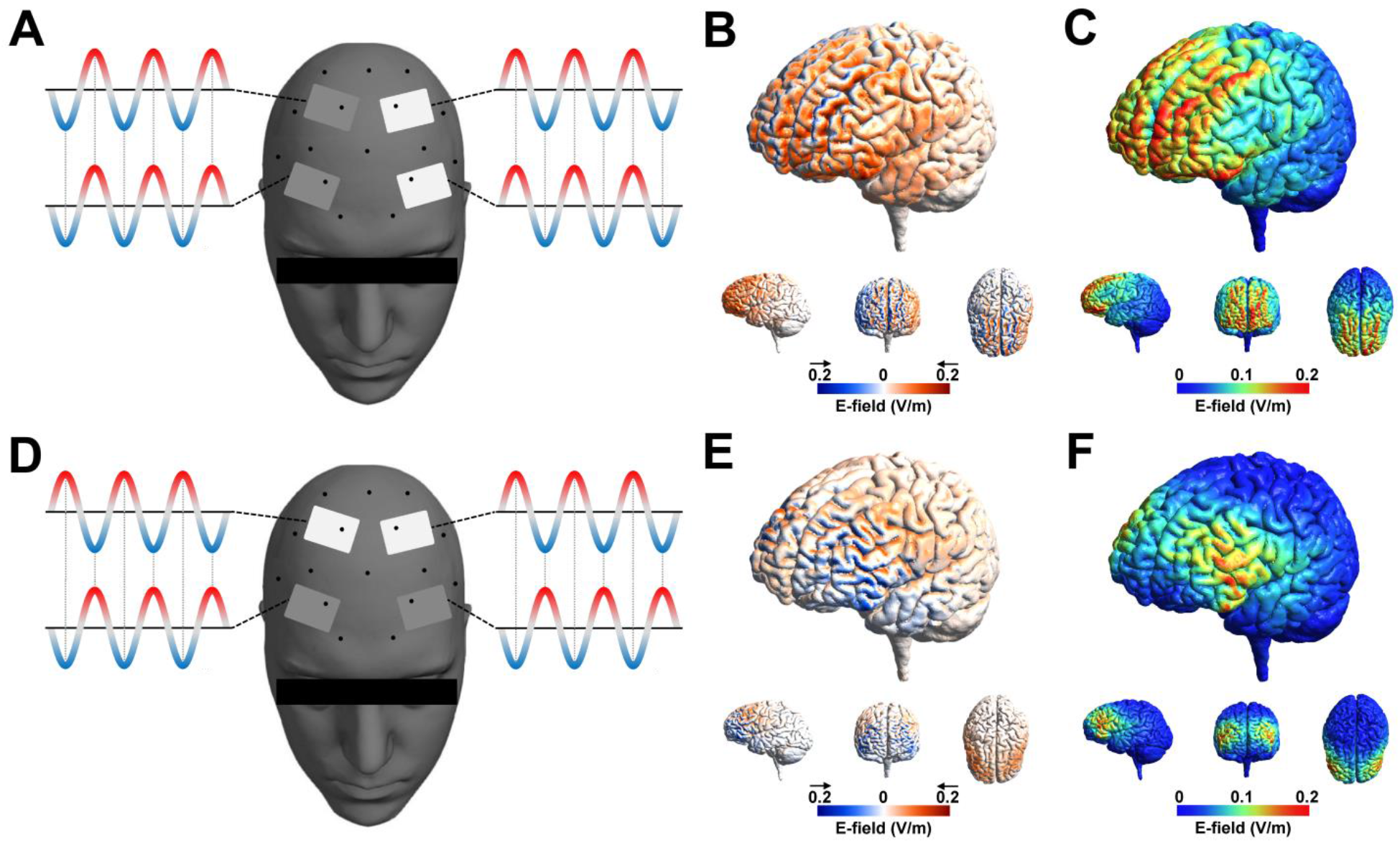
tACS montages and induced electric fields. A) tACS_TRANS_ set up with currents running between AF3 and AF4, as well as Fc1 and Fc2. B) Current flow directionality: tACS_TRANS_ yielded a trans-hemispheric, left-right, current flow. C) Electric field distribution of tACS_TRANS_. D) tACS_INTRA_ set up with currents running between AF3 and Fc1, as well as AF4 and Fc2. E) Current flow directionality: tACS_INTRA_ yielded an intra-hemispheric, posterior-anterior, current flow. F) Electric field distribution of tACS_INTRA_.

Sinusoidal stimulation waveforms were applied at a frequency of 5 Hz with a stimulation intensity of 1 mA peak-to-peak (current density per electrode: 0.067 mA/cm). In all conditions, there was a ramp-up phase of 20 seconds. Stimulation was applied for the entire duration of the task (approximately 30 minutes; total charge: 0.121 C/cm^2^). Sham tACS was applied in the sham condition: after ramp-up, stimulation was applied for 20 seconds, followed by a ramp-down. Current flow direction of sham tACS was counterbalanced. The impedance of the stimulation electrodes was kept below 10 kΩ. After the end of each session, participants reported on whether they had perceived phosphenes or other sensations related to tACS.

### Finite element method (FEM) modelling

SimNIBS 3.2. was used to simulate electric field distributions generated by tACS_INTRA_ and tACS_TRANS_ in a standard head model provided by the SimNIBS software [40]. Default conductivity values were used: skin, fat, and muscles as a single soft tissue (σ = 0.465 S/m), bone (σ = 0.01 S/m), eyes (σ = 0.5 S/m), CSF (σ = 1.654 S/m), grey matter (σ = 0.275 S/m), and white matter (σ = 0.126 S/m). After analysis of the behavioral and EEG results, the commonalities of the electric fields induced by tACS_INTRA_ and tACS_TRANS_ were explored. An averaged electric field distribution of both montages was calculated, as well as the overlapping regions where both electric fields exceeded 0.1 V/m.

### Sensation questionnaires

After each session participants rated the discomfort of sham and active tACS on a five-point scale (adapted from [41]). Perceived discomfort was rated as ‘none’, ‘mild’, ‘moderate’, ‘considerable’ and ‘strong’. Participants rated the following sensations: itching, pain, burning, heat, pinching, metal taste, fatigue, phosphene perception and other. Also, participants were asked how long the sensation lasted with the following possible responses: ‘It stopped quickly’, ‘It stopped half-way’, ‘It lasted until the end’. After completion of the third session, participants were asked to guess during which session placebo (sham) stimulation was applied and how confident they were about their guess.

### Statistical analysis

All statistical analysis were performed using JASP version 0.12.2. Effects of tACS condition (tACS_TRANS_, tACS_INTRA_ and sham) on UV, OE, EF and frontal theta asymmetry shift were investigated by repeated-measures ANOVAs. Significant effects were followed by paired samples t-tests. Reaction time was investigated with a repeated-measures ANOVA with tACS condition and round (first and last round) as independent variables. Theta power was investigated with a repeated measures ANOVA with tACS condition and electrode position (F3 and F4) as independent variables. Partial eta-squared (η_p_^2^) was used as effect size measure for ANOVAs and Cohen’s d was used as effect size for t-test comparisons. Averages are reported ± standard error of mean.

Relationship between behavioral measures (OE, EF and UV) was investigated using Pearson’s correlation coefficient. Additionally, sham-controlled effects of tACS_TRANS_ (tACS_TRANS_ -sham) were correlated to sham-controlled effects of tACS_INTRA_ (tACS_INTRA_ – sham) for each dependent variable, using Pearson’s correlation coefficient.

## Results

No adverse events were reported. Discomfort induced by tACS was rated between ‘none’ and ‘mild’ for all sensations (itching, pain, burning, heat, pinching, metal taste, fatigue and other), with no difference between sham and active tACS (F2,17) = 1.64, p = 0.17). Furthermore, participants reported that any sensation dissipated within the first two minutes in all conditions. None of the participants reported perceiving phosphenes in any stimulation condition. Seven out of 18 participants correctly identified the sham condition, which was not different from chance level χ^2^(1) = 0.25, p = .617.

### Relationship between risk parameters

Correlation analysis of risk parameters UV, OE, and EF revealed a significant negative relationship between OE and EF (r = -0.718, p < 0.001). As expected, this suggests that more exploratory behavior relates to higher general risk-taking behavior. Interestingly, UV was not significantly correlated to OE (r = 0.316, p = 0.201) or EF (r = -0.073, p = 0.774), suggesting that perceived uncertainty is unrelated to general risk taking in this task on a group level. This emphasizes that uncertainty effects on risk taking behavior can differ inter-individually.

### tACS effect on uncertainty

In the last round before stopping, participants reacted significantly slower compared to the first round of a trial (F(1,17) = 34.11, p < 0.001, η_p_^2^ = 0.667; Figure 3A). This increase in time to make a decision reflects perceived uncertainty before stopping and was quantified by the UV (reaction time last round – first round). A repeated-measures ANOVA revealed that UV differed significantly between the stimulation conditions (tACS_INTRA_, tACS_TRANS_, sham), F(2,34) = 3.72, p = 0.034, η_p_^2^ = 0.180 (Figure 3B). Post-hoc t-tests revealed a significant difference between tACS_INTRA_ (133.28 ± 26.37) and sham (90.94 ± 18.39; t(17) = 2.19, p = 0.042, Cohen’s d = 0.52), as well as between tACS_TRANS_ (136.64 ± 23.32) and sham (t(17) = 3.27, p = 0.004, Cohen’s d = 0.77). However, no significant difference was observed between tACS_INTRA_ and tACS_TRANS_ (t(17) = 0.15, p = 0.880, Cohen’s d = 0.04). These results suggest that subjective uncertainty increased when theta tACS was applied, with both montages yielding comparable effects.

**Figure 3.**
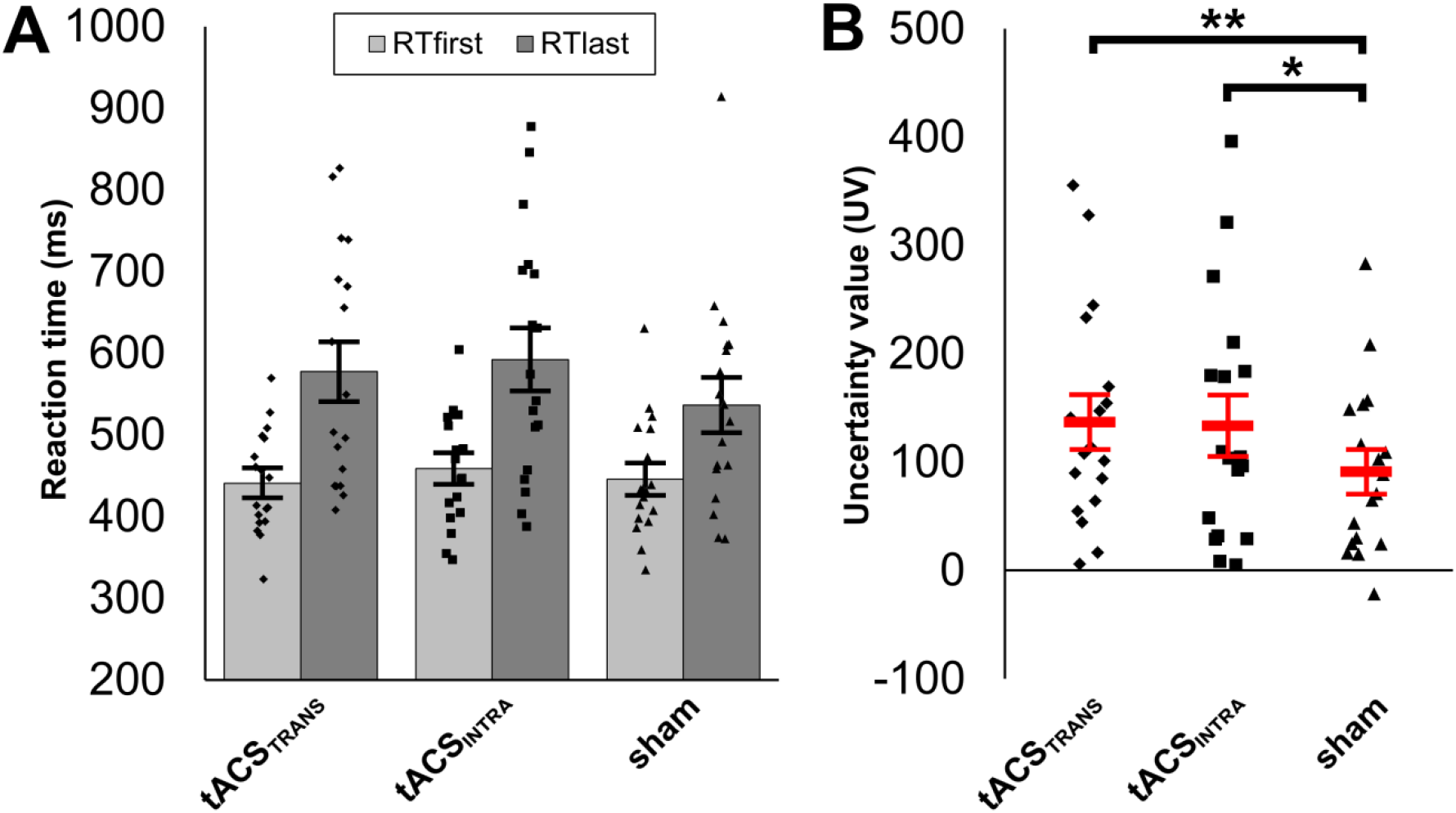
A) Individual and average reaction times during first (RTfirst) and last (RTlast) round of a trial per condition. B) Individual and average uncertainty values (UV; that is RTlast – RT first) for each tACS condition. Error bars representing standard error of mean. *p<0.05, **p<0.01.

### tACS effect on exploratory and general risk-taking behavior

Besides in terms of uncertainty, risk-taking behavior can be described in relation to ones openness to explore various outcome ranges, as well as general bias towards higher rewards. The EF and OE parameters were investigated as a proxy for these two aspects of risk behavior, respectively (Figure 4A,B,C). Repeated-measures ANOVA showed that OE did not significantly differ between the stimulation conditions (F(2,34) = 0.35, p = 0.708, η_p_^2^ = 0.020; Figure 4D). Furthermore, no significant effect of tACS condition on EF was found (F(2,34) = 1.23, p = 0.304, η_p_^2^ = 0.068; Figure 4E). This suggests that, on group level, tACS did not affect exploratory and general risk-behavior.

**Figure 4.**
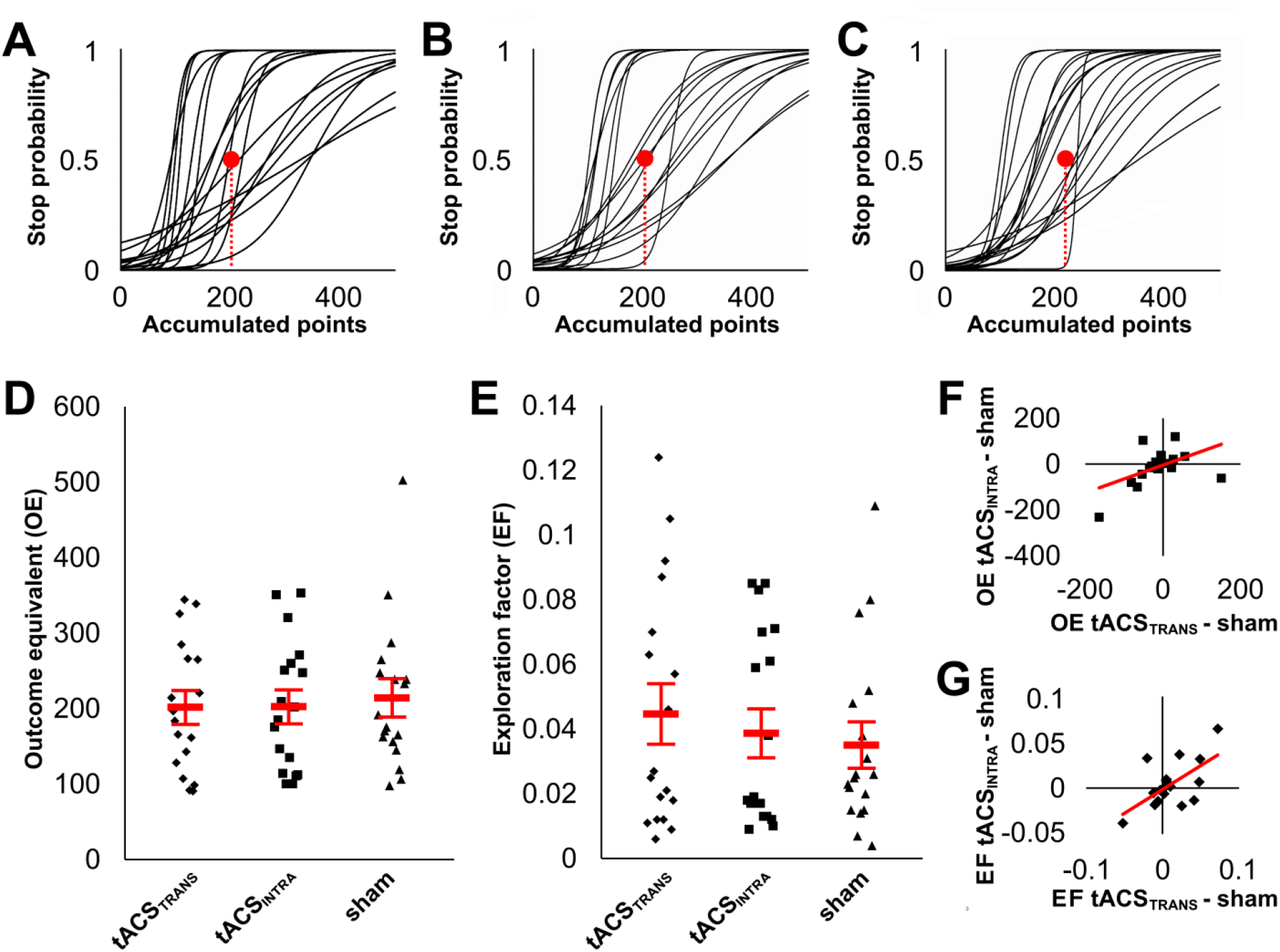
Logistic curves fitted to the stop probability distribution of A) tACS_TRANS_ condition, B) tACS_INTRA_ condition, and C) sham condition. D) Individual and average outcome equivalent (OE) representing general risk-taking for each condition. E) Individual and average exploration factor (EF), representing willingness to explore a risk environment as opposed to exploiting safe rewards. F) Scatterplot representing the correlation between the sham-controlled effect of tACS_TRANS_ (tACS_TRANS_ – sham), compared the sham-controlled effect of tACS_INTRA_ (tACS_INTRA_ – sham) on OE. G) Scatterplot representing the correlation between the sham-controlled effect of tACS_TRANS_ and tACS_INTRA_ on EF. Error bars representing standard error of mean.

Since risk taking is prone to inter-individual variability, effects of tACS_INTRA_ and tACS_TRANS_ were correlated to explore whether behavioral changes on individual data points.. The sham-controlled CE effect of tACS_INTRA_ was significantly positively correlated with the sham-controlled CE effect of tACS_TRANS_ (r = 0.498, p = 0.036; Figure 4F). Similarly, a significant positive correlation on GR was observed between tACS_INTRA_ and tACS_TRANS_ (r = 0.616, p = 0.006; Figure 4G). This implies that participants who increased risk taking and exploratory behavior during tACS_INTRA_ showed a similar effect during tACS_TRANS_ compared to sham, and vice versa. That is, although the directionality of the tACS effect differed between participants, the effect of both montages was comparable.

### tACS effect on frontal theta symmetry

Previous studies have suggested that frontal theta power asymmetry may relate to increased risk-taking behavior. Therefore, we investigated the effect of theta tACS on posttest – pretest changes in frontal theta asymmetry. A repeated measures ANOVA comparing the three stimulation conditions (tACS_INTRA_, tACS_TRANS_, sham) showed a significant effect on theta asymmetry (F(2,34) = 4.01, p = 0.027, η_p_^2^ = 0.191). Post-hoc t-test showed a significant difference between tACS_INTRA_ (0.09 ± 0.02) and sham (−0.01 ± 0.03; t(17) = 3.15, p = 0.006, Cohen’s d = 0.74). No difference was observed between tACS_TRANS_ (0.06 ± 0.04) and sham (t(17) = 1.611, p = 0.126, Cohen’s d = 0.38) and between tACS_INTRA_ and tACS_TRANS_ (t(17) = 0.89, p = 0.388, Cohen’s d = 0.21). We hypothesized that tACS_TRANS_ would induce theta asymmetry, whereas tACS_INTRA_ would promote a balance between hemispheres. Contrary to these expectations, results showed that tACS_INTRA_ induces a right frontal-biased asymmetry.

Moreover, absence of a difference between both tACS montages suggests a similar effect on theta asymmetry. A significant positive correlation between tACS_INTRA_ and tACS_TRANS_ (controlled for sham) effects on frontal theta asymmetry provided additional observation for this observation (r = 0.727, p < 0.001).

Interestingly, investigating the tACS-related change in frontal theta power separately for electrodes F3 and F4 revealed no significant effect of tACS condition (F(2,34) = 2.51, p = 0.096, η_p_^2^ = 0.129; Figure 5B). Furthermore, theta power change did not differ between electrodes (F(1,17) = 0.40, p = 0.534, η_p_^2^ = 0.023). Although a significant tACS*electrode interaction was observed F(2,34) = 3.49, p = 0.042, η_p_^2^ = 0.170), none of the post-hoc t-test comparison were significant (p > 0.06; Figure 5B). In sum, these results suggest changes in frontal theta symmetry are not associated with a tACS-related increase or decrease on theta power. Investigation of delta, alpha and beta power revealed no significant main or interaction effects (p > 0.07).

**Figure 5.**
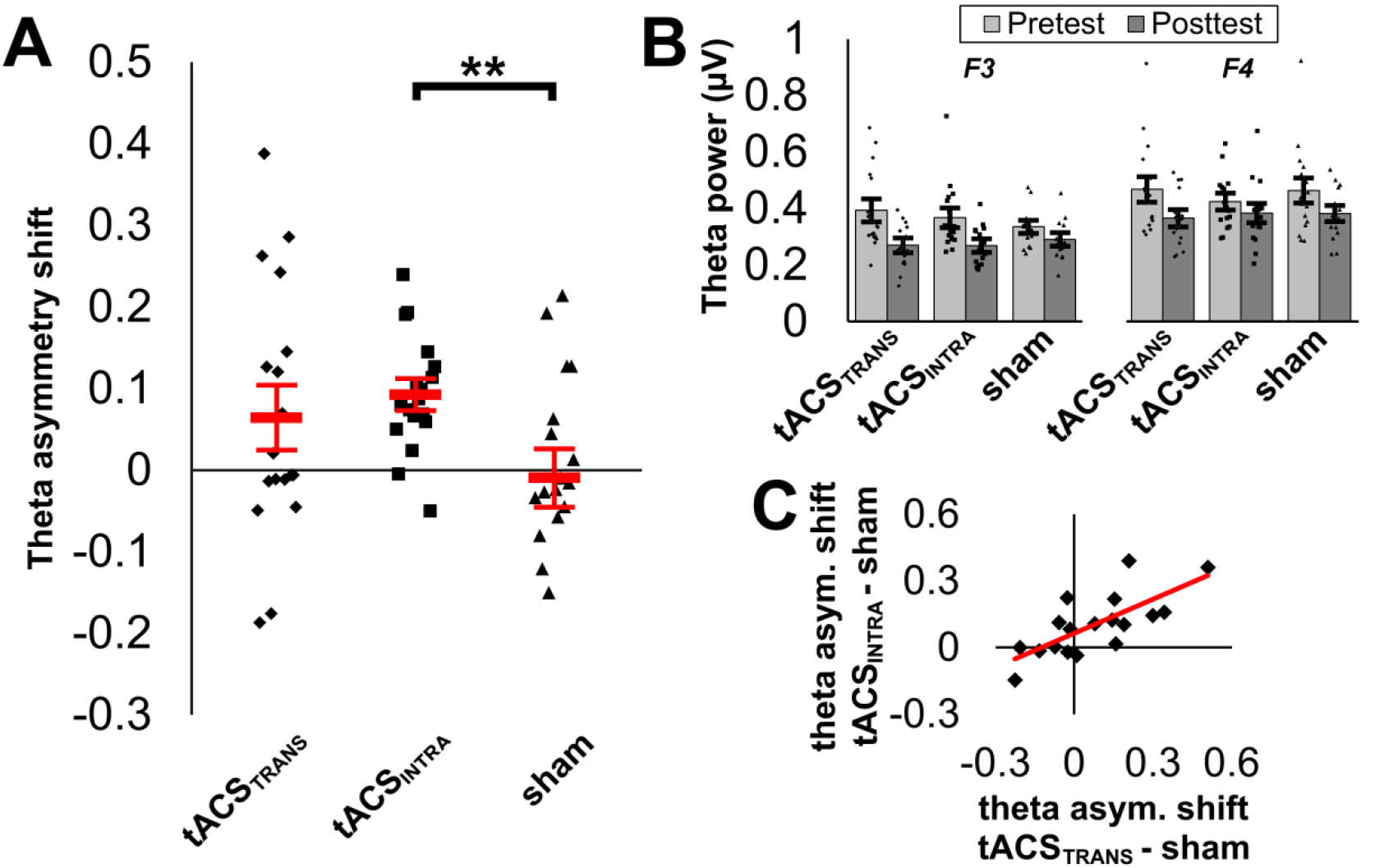
A) Individual and average posttest – pretest difference (shift) in frontal theta asymmetry (spectral activity of F4 – F3 / F4 + F3) for each tACS condition. Positive values indicate a shift towards a right hemispheric bias, whereas negative values indicate a shift towards a left hemispheric bias. B) Pretest and posttest theta power (µV) for each tACS condition in electrodes F3 and F4. C) Scatterplot showing the correlation between the sham-controlled effect of tACS_TRANS_ and the sham-controlled effect of tACS_INTRA_ on the frontal theta asymmetry shift. **p<0.01.

Similar to the behavioral outcomes, the change in theta power was significantly correlated between sham-controlled tACS_INTRA_ and tACS_TRANS._in both electrodes F3 (r = 0.808, p < 0.001) and F4 (r = 0.569, p = 0.014). These results provide further evidence that although the tACS effect varies between participants, both frontal montages induce a similar effect. Importantly, no significant correlations between montages in either electrode were found for delta, alpha and beta power (p > 0.28).

### Post analysis electric field modeling

Despite differences in electric current direction, effects on risk taking (OE and EF) and frontal theta power (asymmetry as well as F3 and F4 individually) were positively correlated between tACS_INTRA_ and tACS_TRANS_. We speculated that observed effects may be driven by commonalities in affected regions between two montages. Therefore, we explored the shared targeted regions of both montages by overlaying electric field distributions. First, the averaged induced electric field was calculated (Figure 6A), followed by an investigation of the overlapping areas of the two montages where both the tACS_INTRA_ and tACS_TRANS_ electric fields were larger than 0.1 V/m (Figure 6B). Average and overlap simulation showed two regions corresponding to the dorsolateral prefrontal cortex (MNI coordinates left:-35 48 30, right: 42 45 28) and the inferior frontal gyrus (MNI coordinates left: -47 41 -9, right: 55, 41, -7).

**Figure 6.**
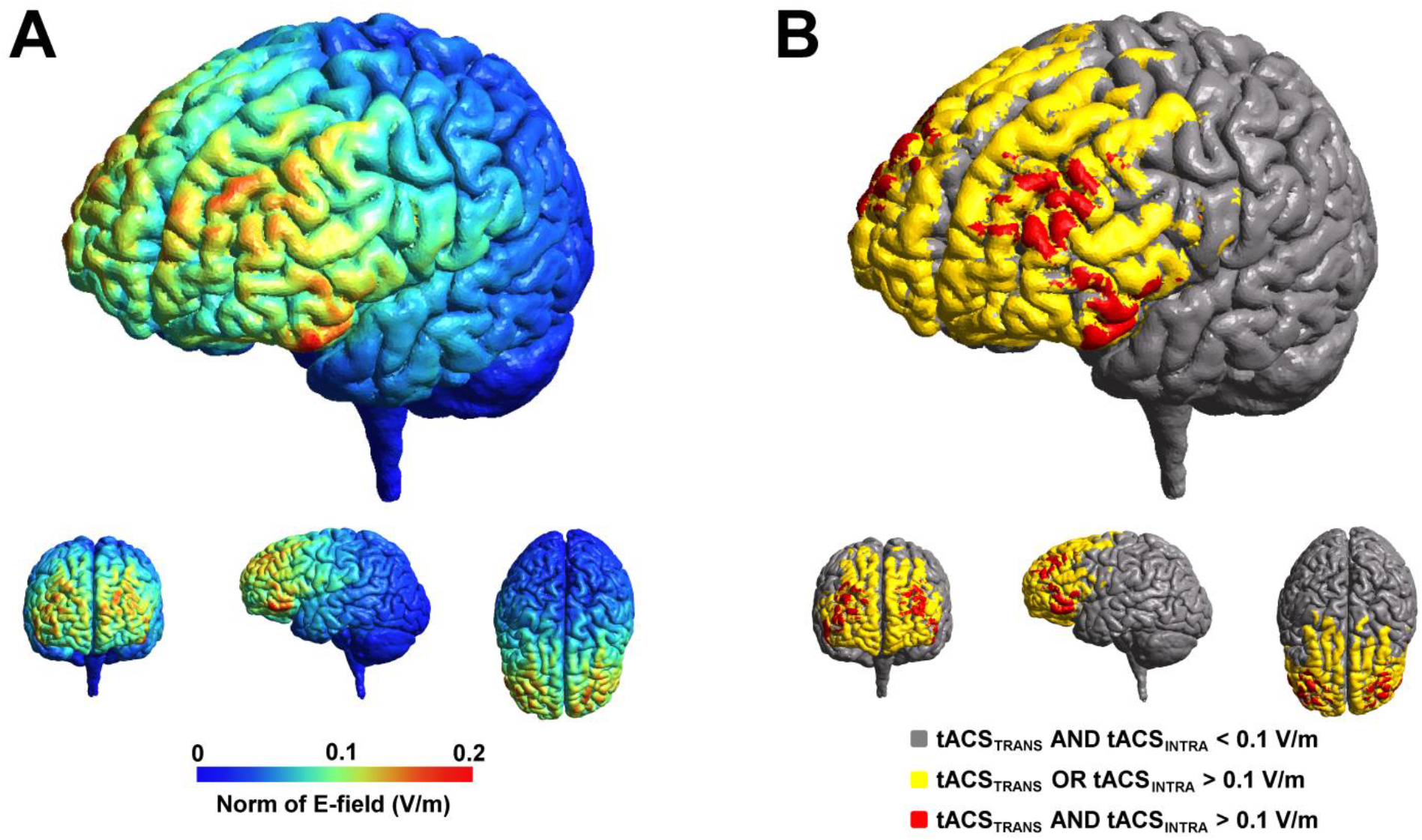
A) Averaged electric field distributions of tACS_INTRA_ and tACS_TACS_. B) Overlapping areas of both tACS montages. Regions where electric fields were larger than 0.1 V/m in both montages shown in red.

## Discussion

In the present study, we found that frontal theta tACS is related to an increase in perceived uncertainty, compared to sham, during a sequential decision-making task with escalating risk. Furthermore, we found that frontal tACS may induce frontal theta asymmetry, resulting in a right-hemispheric dominance. Interestingly, effects on uncertainty and frontal asymmetry were independent of whether applied currents traveled within or between hemispheres. The observed increase in uncertainty was unrelated to other aspects of risk taking that were examined. Consequently, no effect of tACS on exploration behavior and general risk-taking was observed. Despite the absence of a group effect on these variables, individual effects were similar for within-and between-hemispheric montages, providing further evidence suggesting that current flow direction was not crucial to explain the observed effects.

Our observation that theta tACS affects uncertainty during decision-making is in line with previous EEG findings. Uncertainty and the reduction thereof is thought play a crucial role in performance monitoring [42]. Performance monitoring describes a cognitive process of optimizing behavior by reducing prediction errors, striving towards a stable and predictable environment [17, 21-26]. Indeed, substantial evidence points towards a relation between frontal theta oscillations, prediction error processes and uncertainty reduction (for a review see [43]). Furthermore, Reinhart & Woodman [44] showed that modulation of theta-mediated electrocortical error signals by frontal transcranial direct current stimulation (tDCS) relates to behavioral adaptation. Thus, frontal theta activity and uncertainty underlie updating of internal prediction models that may inform behavior in response to rewards and punishments.

Interestingly, although tACS was related to an increase in uncertainty, we observed no group effect on exploratory (EF) and general risk behavior (OE). Uncertainty can be reduced by opting for a low risk approach of exploiting known outcomes in some situations. However alternatively, exploratory behavior may also promote uncertainty reduction by obtaining more contextual information on potential positive or negative outcomes [3, 45]. Thus, whether uncertainty dictates if one needs to explore or exploit in risky situations depends on environmental and individual factors [3]. Participant-specific (individual) features, such as reward sensitivity, loss aversion, cognitive capacity, emotional state and frontal cortex activation have an impact on subjective uncertainty [6,7,46-51]. In addition, task-related (environmental) features, such as reward probability, reward value, cost of information and predictability determine whether exploration or exploitation will be preferred [52, 53]. Interestingly, whether uncertainty relates to exploratory behavior may be associated with activity in the rostrolateral PFC [7]. Consequently, whether uncertainty correlates with risk-taking depends on the specific behavioral task. Individual and task-related factors may provide an explanation for why a tACS-related change in uncertainty did not translate into a difference in risk behavior in our study. In accordance, several studies investigating the effect of frontal tDCS on risk-taking did observe changes in perceived uncertainty, without effects on risk-taking behavior [54, 55]. In sum, individual and environmental variability may explain why previous studies, using different behavioral tasks, did find theta tACS-related effects on overall risk behavior [19, 27], whereas in the present study only uncertainty was affected.

Post-to pre-test comparison of resting state EEG showed a change to a right-hemispheric bias in frontal theta asymmetry. Previous studies suggested that modulating frontal theta asymmetry with tACS may modulate risk-taking behavior [19, 27]. Sela et al. [19] showed that right frontal tACS may increase risk taking, whereas Dantas et al. [27] showed that left frontal tACS may decrease risk-raking. Since we observed no tACS-related effects on risk behavior and EEG was recorded before and after the task, our findings cannot provide direct evidence for an associated between risk-taking and frontal asymmetry. Notwithstanding this limitation, the observation that theta tACS might modulate oscillatory symmetry is interesting, as frontal asymmetries have been described as neural correlates to a variety of affective and cognitive functions [56-58]. Contrary to our expectations, no difference in frontal asymmetry change was observed between tACS_INTRA_ and tACS_TRANS_. Although we speculated that the direction of current flow may differentially affect hemispheric asymmetry, our results provide no evidence for this hypothesis. The observation of similar tACS-related electrocortical effects may explain why findings on subjective uncertainty, exploration, and general risk-taking were similar for both tACS conditions.

Behavioral effects on UV, OE and EF, as well as changes in resting-state theta power and asymmetry were positively correlated between tACS_TRANS_ and tACS_INTRA_. These observations suggest that, although the response to tACS may vary inter-individually, effects were similar for both montages. One potential explanation is that the neuromodulation-induced changes were driven by targeted regions that both montages had in common and not by current flow direction. That is, regions where induced electric fields overlap. Therefore, we performed an electric field simulation that highlights these commonalities. Interestingly, electric field simulation suggested that both the DLPFC and IFG were primarily targeted by both tACS set ups. Coincidentally, these areas were identified by Meder et al. [8] to reflect individual differences in performance on the same risk-taking task. Since the DLPFC has been related to exploratory behavior and the IFG to exploitation behavior, a balance between both areas may arguably be associated with individual risk-taking tendencies.

It should be noted that observations in these post-hoc simulations are not conclusive and should be interpreted with caution. As such, alternative explanations for our findings have to be considered. For instance, (trans)cutaneous effects of tACS may contribute to observed changes in behavior [59, 60]. Since in this study both tACS conditions resulted in similar outcomes, it could be argued that the present findings may be explained, at least in part, by cutaneous effect. However, sensation questionnaires showed that participants did have different experiences during active tACS compared to sham. Furthermore, similarity of effects on resting state EEG were specific to the theta frequency. Therefore, we feel that cutaneous effects are not sufficient to explain our findings, but can also not be fully excluded.

Another point to consider is that tACS-induced electric fields were relatively small with an approximate peak of 0.2 V/m. Although such current intensity over prefrontal cortex may induce changes in cognitive performance [28,29, 61-64], electrical brain stimulation effects are subject to a large amount of inter-and intra-individual variability [65, 66]. Furthermore, it has been suggested that higher current intensities can yield more consistent effects [32, 67]. However, when interpreting electric field simulations it should also be considered that the standard model was used. Since induced electric fields can differ significantly between participants depending on head morphology, the actual induced currents in the present study might differ [68].

In conclusion, this study shows that frontal theta tACS can increase subjective uncertainty in a risk-taking task. Although previous studies have shown that an increase in perceived uncertainty may spark either exploration or exploitation in risky environments, such an association was absent in the sequential gambling task that we employed. As such, no effect of tACS on exploratory behavior or general risk-taking was observed. Furthermore, it was found that frontal theta tACS may induce a shift in theta asymmetry towards a right-hemispheric bias, independent of electric current flow direction. Instead, our findings may be explained by the overlap between tACS-affected cortical region of both montages used. Altogether, our findings provide the first insight into how tACS may influence specific exploration-exploitation dimensions of risk-related decision-making.

